# Brief report: Reclassifying SCLC-Y as SMARCA4 deficient malignancies - resolving the controversy

**DOI:** 10.1101/2022.10.09.511505

**Authors:** Jin Ng, Ling Cai, Luc Girard, Owen W.J. Prall, Neeha Rajan, Christine Khoo, Ahida Batrouney, Ariena Kersbergen, Michael Christie, John D. Minna, Marian L. Burr, Kate D. Sutherland

## Abstract

**Introduction:** The classification of small cell lung cancer (SCLC) into distinct molecular subtypes defined by ASCL1, NEUROD1, POU2F3 or YAP1 expression, paves the way for the development of targeted therapeutics. However, the existence of a distinct YAP1-expressing SCLC subtype remains controversial. Here we have undertaken a detailed molecular and histological characterisation of YAP1 expressing SCLC-Y to understand the biology of this proposed subtype.

**Methods:** The mutational landscape of human SCLC cell lines was interrogated to identify pathogenic genomic alterations unique to SCLC-Y. Xenograft tumours generated from cell lines representing the molecular subtypes of SCLC (SCLC-A, -N, -P and -Y) were evaluated by a panel of pathologists. Diagnoses were validated by transcriptomic analysis of primary tumour and human cell line datasets.

**Results:** Unexpectedly, pathogenic mutations in SMARCA4 were identified in six of eight SCLC-Y cell lines and correlated with reduced SMARCA4 mRNA and protein expression. Pathologist evaluations revealed that SMARCA4-deficient SCLC-Y tumours exhibited features consistent with thoracic SMARCA4-deficient undifferentiated tumours (SMARCA4-UT). Similarly, the transcriptional profile SMARCA4-mutant SCLC-Y lines more closely resembled primary SMARCA4-UT, or SMARCA4-deficient non-small cell carcinoma, than SCLC. Combining clinical, pathological, transcriptomic, and genetic data we found little evidence to support a diagnosis of SCLC for any of the YAP1-expressing cell lines originally used to define the SCLC-Y subtype.

**Conclusions:** SCLC-Y cell lines harbour inactivating *SMARCA4* mutations and exhibit characteristics consistent with SMARCA4-deficient malignancies rather than SCLC. Our findings suggest that, unlike ASCL1, NEUROD1 and POU2F3, YAP1 is not a subtype defining transcription factor in SCLC.

## INTRODUCTION

Small cell lung cancer (SCLC) is a highly aggressive form of lung cancer, for which effective treatments are desperately needed. While SCLC has traditionally been treated as a single disease entity, four distinct molecular SCLC subtypes, based on expression of key transcription factors (TFs) ASCL1 (SCLC-A), NEUROD1 (SCLC-N), POU2F3 (SCLC-P) and YAP1 (SCLC-Y), have recently been identified. This seminal advance in the field has created hope for the development of subtype-specific molecularly targeted therapies. Understanding the biology of these different subtypes is therefore paramount^1^.

SCLC-Y (defined by the expression of YAP1) has proven the most difficult of these subtypes to define, in part because it is characterised by low expression of classical neuroendocrine markers synaptophysin, chromogranin A, NCAM1 (CD56) and INSM1. 5-10% of SCLC may show low or absent neuroendocrine marker expression^2^ and a diagnosis of SCLC can be made in the absence of positive neuroendocrine marker immunohistochemistry if the tumour morphology is characteristic^3^. However, neuroendocrine marker low/negative SCLC presents a diagnostic challenge as many other malignancies that can mimic SCLC histopathology must be excluded. While YAP1 expressing, neuroendocrine low SCLC cells have been detected in patient-derived xenografts, genetically engineered mouse (GEM) models and some patient cohorts^4–6^, other studies have failed to identify a distinct YAP1-positive SCLC subtype^1,7^. Together, these observations have fuelled the controversy around the existence of SCLC-Y as a *bona fide* SCLC subtype^1,6,7^.

The initial identification and classification of the SCLC-Y subtype was based predominantly on transcriptome profiling of human SCLC cell lines^8,9^. These lines provide valuable, well characterised SCLC disease models that are widely used in the field^10,11^. To better understand the biology of SCLC-Y, we therefore undertook a comprehensive molecular and histological characterisation of the tumour lines used to define this proposed subtype of SCLC.

## RESULTS

### Pathogenic *SMARCA4* mutations are enriched in SCLC-Y

SCLC almost universally harbours mutations of both *TP53* and *RB1*, however SCLC-Y frequently lack *RB1* mutations^9^. We therefore interrogated the mutational landscape of 50 SCLC cell lines used to classify the SCLC-Y subtype (Cancer Cell Line Encyclopaedia, CCLE^12^) and identified 26 frequently mutated genes unique to SCLC-Y cell lines (**Fig. 1*A*** and **Supplementary Table 1**). This dataset included all eight YAP1 expressing SCLC lines previously included in the classification of the SCLC-Y lineage^8^. Notably, mutations in *SMARCA4* (also known as *BRG1*), which encodes an ATPase subunit of SWI/SNF chromatin-remodelling complexes^13^, were observed in six out of eight SCLC-Y lines and found mutually exclusive with mutations in *RB1* (**Fig. 1*B* and Supplementary Fig. 1*A***). All cancer cell lines harbouring frame shift or nonsense mutations in *SMARCA4* showed reduced SMARCA4 mRNA and protein abundance (**Fig. 1*C* and Supplementary Fig. 1*B***). All six SMARCA4 mutant SCLC-Y lines also harboured *TP53* mutations, but other mutations previously observed to co-occur with SMARCA4 mutations in NSCLC, such as *STK11, KEAP1* and *KRAS* were not identified (**Supplementary Table 2**). Consistent with the specific enrichment of SMARCA4 mutations in SCLC-Y tumours, *SMARCA4-*-mutant SCLC exhibited significantly lower expression of a previously defined neuroendocrine transcriptomic signature^10^ compared to *SMARCA4* wildtype SCLC (**Fig. 1*D*** and **Supplementary Table 3**).

**Figure 1.**
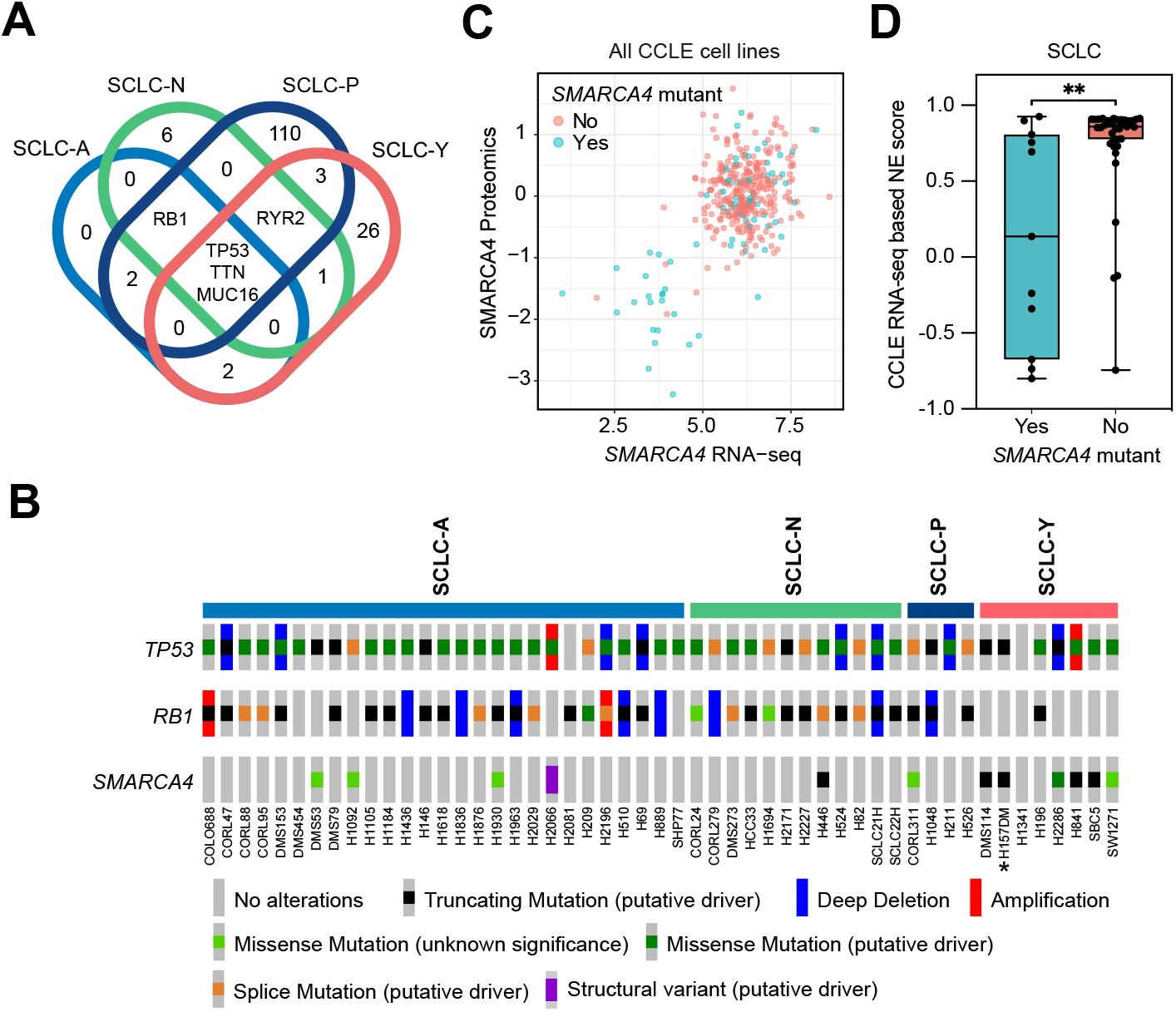
*SMARCA4* mutations are enriched within SCLC-Y cell lines. (***A***) Genetic mutations that are present in at least 50% of samples within each SCLC subtype group were identified. These co-occurring and subtype-specific mutations associated with each SCLC molecular subtype are displayed in the Venn diagram. There are 26 mutations exclusive to SCLC-Y cell lines (see **Supplementary Table 1**). (***B***) Of all 26 mutations identified, *SMARCA4* mutations were present in six of eight SCLC-Y cell lines, together with *TP53* mutations on a *RB1* wildtype background. *H157DM was previously annotated in CCLE as H1339. (***C***) All cell lines in the CCLE (lung and non-lung cancer cell lines) that had complete proteomic and RNA-seq profiles were interrogated for the correlation between SMARCA4 protein and mRNA in cell lines that had mutant and wildtype *SMARCA4*. There is a positive correlation (Pearson correlation = 0.53, p=1.5e-27) between low *SMARCA4* mRNA and loss of SMARCA4 protein. (***D***) NE scoring was performed using a previously published 50-gene transcriptomic signature^10^ on SCLC cell lines in the CCLE. SCLC cell lines with *SMARCA4* mutations have a significantly lower NE-score compared to wildtype *SMARCA4* SCLC cell lines (Mann-Whitney test, **p=0.0068).

### Histopathological classification of SMARCA4-deficient SCLC-Y

Inactivating mutations in *SMARCA4* are characteristic of small cell carcinoma of the ovary, hypercalcaemic type (SCCOHT) and can also occur in malignant rhabdoid tumours. Within the thorax, SMARCA4 loss is seen in two different settings; either in non-small cell carcinoma (NSCLC) harbouring SMARCA4 mutations^14^, or in thoracic SMARCA4-deficient undifferentiated tumours (previously named SMARCA4-deficient thoracic sarcoma)^15^. While SMARCA4-deficient NSCLC (frequently adenocarcinomas) may display histological features of de-differentiation including loss of TTF-1 expression and a solid growth pattern; thoracic SMARCA4-deficient undifferentiated tumours (SMARCA4-UT) show undifferentiated round cell or rhabdoid features and reduced expression of epithelial markers such as cytokeratins and Claudin-4. SMARCA4 mutations have not previously been reported in SCLC. Therefore, to better define the identity of SMARCA4-mutant SCLC-Y tumours, we established a panel of SCLC cell line xenograft models derived from all SCLC transcriptional subtypes, including 3 SMARCA4-mutant SCLC-Y lines. As SCLC diagnosis is primarily based on morphological assessment of the tumour^16^, five pathologists who routinely report thoracic pathology were asked to independently evaluate the de-identified H&E slides and indicate whether the morphology was consistent with SCLC (**Fig. 2*A***). Although there was some variability in individual pathologist assessments for SCLC-A, -N and -P tumours, there was near complete consensus that the H&E appearances of four of five SCLC-Y tumours were not consistent with SCLC (**Fig. 2*B*** and **Supplementary Fig. 2*A&B***). Pathologists were then provided with an immunohistochemical panel (**Supplementary Table 4**) and asked to provide an updated diagnosis (**Fig. 2*C***). Immunohistochemistry for SMARCA4 confirmed that the three SCLC-Y tumours harbouring SMARCA4 mutations were SMARCA4-deficient, whereas SMARCA4 expression was retained in all other tumours (**Fig. 2*D*** and **Supplementary Table 4**). All three of these SMARCA4-deficient SCLC-Y tumours were favoured by pathologists to represent either SMARCA4-UT or SMARCA4-deficient NSCLC rather than SCLC^3^ (**Fig. 2*C***). In contrast to other SCLC subtypes, these SCLC-Y tumours showed expression of RB1, weak cytokeratin staining and isolated expression of synaptophysin in the absence of expression of other neuroendocrine markers INSM1 and NCAM1/CD56 (**Fig. 2*D*** and **Supplementary Fig. 3*A***). This immunohistochemical profile would be highly unusual for SCLC but is characteristic of thoracic SMARCA4-UT^17^. We observed high concordance between neuroendocrine marker mRNA and protein expression (**Supplementary Fig. 3*B***). Of the two SMARCA4 proficient SCLC-Y lines, only one was considered morphologically and immunophenotypically consistent with SCLC (H1341) (**Fig. 2*B&C*** and **Supplementary Fig. 3*B***). Conversely, all five pathologists independently designated H196 to be a sarcomatoid (spindle cell) malignancy (**Fig. 2*B&C***), with a differential diagnosis including sarcomatoid carcinoma, sarcoma, malignant mesothelioma and melanoma.

**Figure 2.**
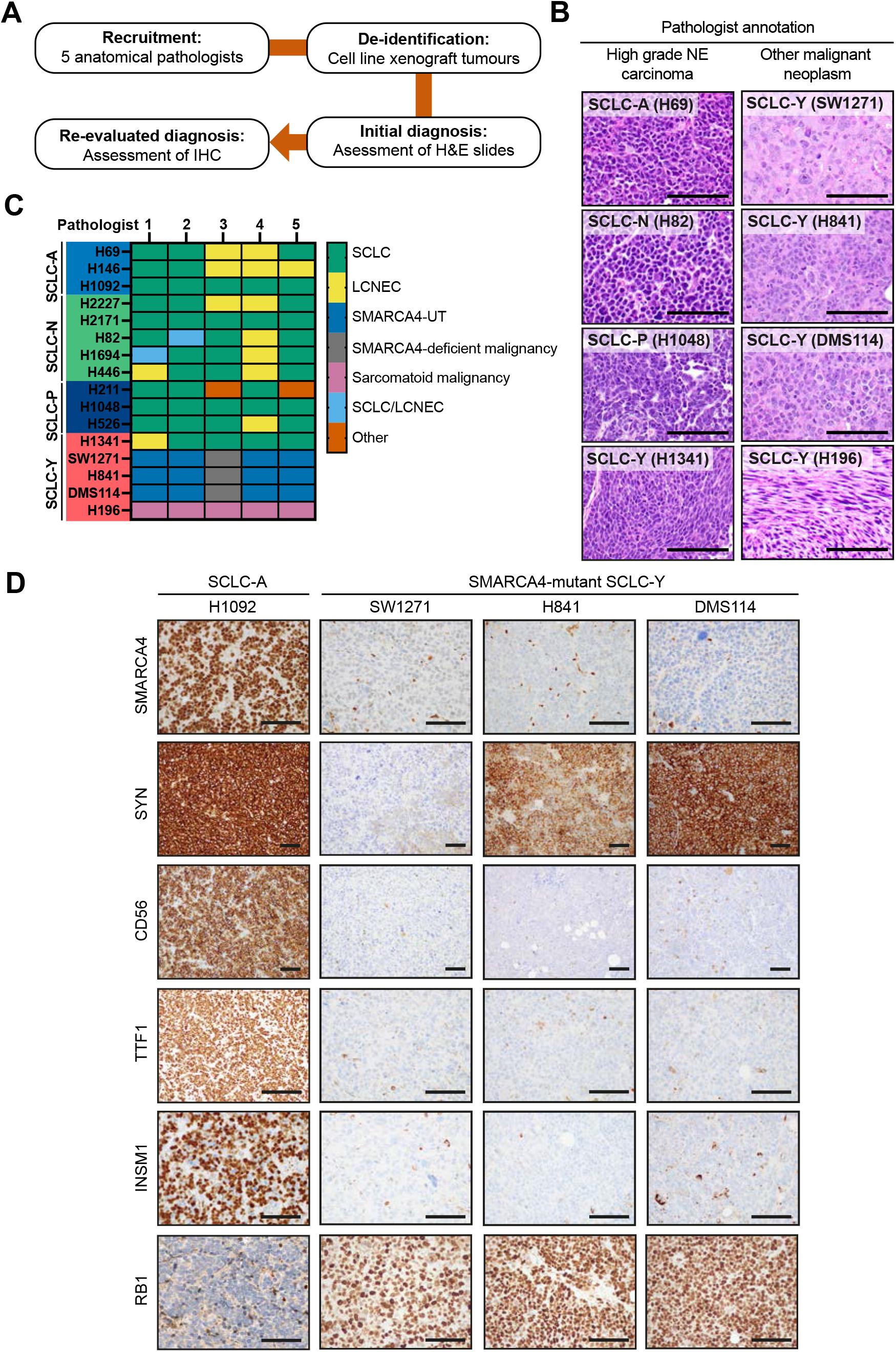
SMARCA4-deficient SCLC cell lines are morphologically and immunophenotypically similar to SMARCA4-UT. (***A***) Methodology outline for assessing SMARCA4-deficient SCLC cell lines by a panel of five anatomical pathologists. Pathologists were provided with H&E staining of cell line xenografts representing each of the four SCLC subtypes and asked to provide an initial diagnosis (**Supplementary Fig. 2*A***). Following the initial assessment, the pathologists were then provided with representative images of IHC staining plus a table summarising the IHC staining profile of each cell line (**Supplementary Table 3**) and asked to provide an updated diagnosis. (***B***) Representative H&E images of cell line xenografts diagnosed as high-grade NE carcinoma (SCLC or LCNEC) and other malignant neoplasm (Not SCLC) by the panel of pathologists. Scale bar = 100 μm. (***C***) Heatmap of re-evaluated diagnosis of SCLC cell line xenografts (row) and each pathologists’ classification (column). There is a consensus among pathologists for diagnosing high grade NE carcinoma (SCLC or LCNEC) among the SCLC-A, SCLC-N and SCLC-P cell line xenografts. Conversely, only one SCLC-Y cell line, H1341, fit the diagnostic criteria for SCLC. Abbreviations: SCLC = Small cell lung cancer; LCNEC = Large cell neuroendocrine cancer; SMARCA4-UT = SMARCA4-deficient undifferentiated tumour; SCLC/LCNEC = Combined SCLC and LCNEC components; Other = Undifferentiated tumour or small blue round cell tumour. (***D***) Representative immunohistochemical images of a NE-high SCLC cell line (H1092) compared to SMARCA4-mutant SCLC cell lines. SMARCA4-mutant SCLC cell lines showed loss of SMARCA4 protein, retained RB1, variable synaptophysin and absent CD56, TTF1 and INSM1 staining. Scale bar = 100 μm.

To further validate the identity of the SCLC-Y lines we reviewed the data in CCLE and compared to independent DNA-sequencing, RNA-sequencing and histological analysis of the same tumour lines undertaken at UT Southwestern Medical Center (UTSW) (**Supplementary Table 5**). This comparison demonstrated high concordance between independent sequencing datasets but revealed that two of the eight previously annotated SCLC-Y lines^8^, were derived from tumours with a primary histological diagnosis of NSCLC rather than SCLC. The CCLE line H157DM (*TP53*^mut^ *RB1*^wt^, *SMARCA4*^truncating^) was previously thought to be the SCLC line H1339 but was more recently identified to be the lung squamous cell carcinoma line H157, an identity that was confirmed on comparison with UTSW data (**Supplementary Table 5**). In addition, H2286 (*TP53*^mut^, *RB1*^wt^, *SMARCA4*^missense^) was derived from a tumour that showed mixed histology and was classified as an adenocarcinoma by UTSW according to an adenocarcinoma-squamous cell carcinoma RNA-seq signature^18^. Thus, in addition to the five SCLC-Y lines directly examined in this study, these findings suggest that an additional two previously designated SCLC-Y lines (H1339/H157DM and H2286) are SMARCA4-mutant NSCLC rather than SCLC.

### SMARCA4-deficient SCLC cell lines have a similar transcriptome to SMARCA4-UT

*SMARCA4* and *SMARCA2* encode mutually exclusive core ATPase subunits of SWI/SNF chromatin remodelling complexes^13,14^. In contrast to SMARCA4-deficient NSCLC in which SMARCA2 is essential for survival^19^, thoracic SMARCA4-UT and SCCOHT typically exhibit concurrent silencing of SMARCA2 expression^17,20^. Disruption of SWI/SNF function in these tumours leads to broad transcriptional dysregulation and remarkably this epigenetic reprogramming appears to be a more dominant driver of phenotype than the tissue of origin. Thus, the transcriptome of thoracic SMARCA4-UT more closely resembles SCCOHT than SMARCA4-deficient NSCLC^15^. To investigate whether the transcriptional profile of SMARCA4-mutant SCLC-Y lines are closely related to SMARCA4-UT or to other SCLC subtypes, we integrated RNA sequencing data from all SCLC, NSCLC and SCCOHT cell lines in CCLE with RNA sequencing data from primary tumours, including primary thoracic SMARCA4-UT, SCCOHT and unclassified thoracic sarcomas^15^. Unsupervised hierarchical clustering using a previously defined set of differentially expressed genes in primary SMARCA4-UT compared to SMARCA4-deficient NSCLC^15^ revealed clustering of three SMARCA4-deficient SCLC-Y lines (H841, DMS114, SBC5) with primary thoracic SMARCA4-UT and SCCOHT (**Fig. 3*A***). Conversely, the other SMARCA4-deficient SCLC-Y lines (SW1271, H2286) clustered with SMARCA4-mutant NSCLC rather than SCLC (**Fig. 3*A***). This observation was replicated through principal component (PC) analysis (**Supplementary Fig. 4*A***). In addition to the SCLC-Y lines, four SMARCA4-mutant NSCLC lines (H522, H2077, H1581, H661) localised to the SMARCA4-UT/SSCOHT cluster (**Fig. 3*A***). The primary tumours from which the H1581 and H661 cell lines were derived, lacked morphological features of differentiated adenocarcinoma or squamous cell carcinoma (**Fig. 3*B***). Furthermore, all the SMARCA4-deficient lung cancer cell lines localising to this SMARCA4-UT cluster showed a gene expression profile characteristic of SMARCA4-UT and distinct from that of SCLC and SMARCA4-mutant NSCLC^17^ (**Fig. 3*A* and Supplementary Fig. 4*A***). This included SMARCA4/SMARCA2 co-deficiency, low Claudin-4, variable expression of stem cell markers SALL4, SOX2 and CD34, synaptophysin expression and absent INSM1 and NCAM1 expression (**Fig. 3*C*** and **Supplementary Fig. 4*B***).

**Figure 3.**
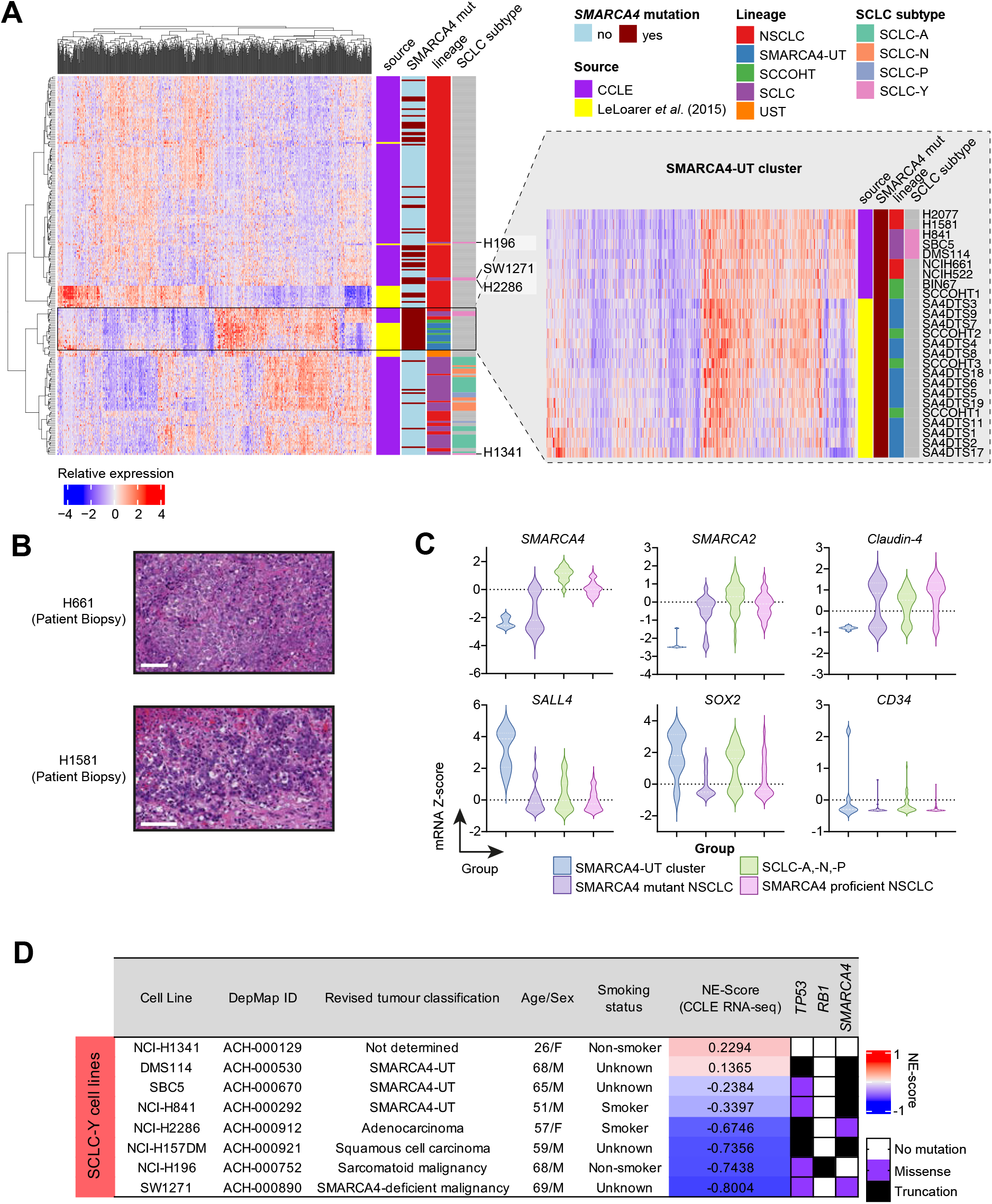
SCLC-Y cell lines with SMARCA4 loss share a similar transcriptome to SMARCA4-UT. (***A***) Unsupervised hierarchical clustering was performed on SCCOHT and lung carcinoma samples (cell lines from the CCLE and bulk-RNA sequenced patient samples from Le Loarer *et al*.^15^). The clustering was based on genes differentially expressed in primary SMARCA4-UT compared to primary SMARCA4-mutant NSCLC identified by Le Loarer *et al* (n=758 genes). SMARCA4-deficient SCLC-Y lines DMS114, H841 and SBC5 clustered with primary SMARCA4-UT and SCCOHT tumours, while SW1271 and H2286 clustered with SMARCA4-deficient NSCLC cell lines. (***Inset***) SMARCA4-UT cluster from panel (***A***) with cell line and sample annotation. Additional lung cancer cell lines identified in the SMARCA4-UT cluster include H661 and H1581, which have been previously classified as NSCLC. (***B***) H&E images of the original patient biopsy used to derive H661 and H1571. Scale bar = 80 μm (***C***) Gene expression of *SMARCA4* and *SMARCA2* together with other clinical markers of SMARCA4-UT (*Claudin-4, SALL4, SOX2* and *CD34*) within cell lines of the SMARCA4-UT cluster, SMARCA4-mutant NSCLC, SCLC-A, -N, -P, and SMARCA4 proficient NSCLC. (***D***) Summary of revised tumour classification for previously designated SCLC-Y lines. H157-DM was previously named H1339 in CCLE.

Taken together, our findings demonstrate that 7 of 8 cancer cell lines originally used to classify the SCLC-Y subtype of SCLC show molecular, morphological and/or transcriptomic features that are inconsistent with a diagnosis of SCLC. One SCLC-Y line (H1341) was pathologically consistent with SCLC, however, the clinical and molecular features of this tumour are unusual for SCLC, as discussed below. Critically, six lines harbour SMARCA4 mutations and display features of either SMARCA4-deficient NSCLC or thoracic SMARCA4-UT.

## DISCUSSION

Understanding SCLC heterogeneity and the distinct biology of the proposed SCLC subtypes is critical for the development of targeted therapies for this devastating disease. Here we shed light on the origins of the controversial SCLC-Y subtype, making the surprising observation that the majority of SCLC-Y cell lines, on which the classification of SCLC-Y was largely based, harbour SMARCA4 mutations. Detailed molecular and histopathological characterisation of these tumours revealed that these tumours show features in keeping with SMARCA4-deficient malignancies rather than SCLC.

Several of the SCLC-Y tumour lines showed close transcriptional similarities with primary thoracic SMARCA4-UT. These tumours showed pathological features consistent with SMARCA4-UT and gene expression profiles consistent with this diagnosis, including loss of both SMARCA4 and SMARCA2 expression and low Claudin-4. In contrast to SCLC-A, -N and -P subtypes, these SMARCA4-deficient SCLC-Y tumours retained RB1 expression and expressed synaptophysin but lacked both NCAM1 (CD56) and INSM1, a profile that is characteristic of SMARCA4-UT.

The classification of SCLC subtypes was based both on primary SCLC samples as well as SCLC cell lines, however, cell lines were particularly enriched within the SCLC-Y category^8^. All of these lines were developed from patients clinically diagnosed as having SCLC, in nearly every case by pathologists recognised as world experts in SCLC diagnosis working as part of a group with extensive experience in SCLC clinical trials (NCI-Navy Medical Oncology Branch)^21^. Nevertheless, these diagnoses were made well before our current knowledge of the key genomic features and immunohistochemical markers characteristic of SCLC and other lung tumours was established and prior to the identification of SMARCA4 deficient malignancies. Thoracic SMARCA4-UT is a recognised mimic of SCLC, and 23% of SMARCA4-UT in a recent series were initially diagnosed as small cell or large cell neuroendocrine carcinoma^17^. Like SCLC, SMARCA4-UT typically occurs in middle-aged smokers and can mimic several histological features of SCLC including small cell morphology, high proliferation index, crush artefact and synaptophysin expression^17^. These features may account for the original SCLC diagnosis, however the morphology and immunohistochemical profile of the SMARCA4-deficient SCLC-Y xenografts was unanimously considered by a panel of pathologists to be consistent with thoracic SMARCA4-UT or SMARCA4-deficient carcinoma as opposed to SCLC. Although we did not identify SMARCA4 mutations in two of the eight SCLC-Y lines, one of these lines was pathologically not consistent with SCLC. The other line was isolated from a cervical tumour deposit in a 26-year-old female non-smoker and lacks mutations of either *TP53* or *RB1*, and so is also unlikely to be SCLC. Interestingly, this tumour has an intragenic deletion in SMARCB1, raising the possibility that this tumour could also represent a SWI/SNF deficient malignancy. Thus, altogether we found little evidence to support a diagnosis of SCLC for any of the SCLC-Y lines tested (**Fig. 3*D***)

The expression of YAP1 is inversely correlated with the expression of neuroendocrine markers in SCLC, and thus ‘classical’ neuroendocrine high SCLC lacks expression of YAP1. In contrast, YAP1 is expressed in NSCLC, including adenocarcinoma, squamous cell carcinoma and a proportion of large cell neuroendocrine carcinomas^7,22^. The expression of YAP1 in multiple lung malignancies therefore complicates the use of YAP1 expression to define a specific subtype of SCLC, particularly given the occurrence of combined tumours in which YAP1 expressing NSCLC may be admixed with SCLC, and the existence of YAP1-positive tumours that can mimic SCLC histologically, such as basaloid squamous cell carcinoma, poorly differentiated adenocarcinoma, high-grade adenoid cystic carcinoma and SMARCA4-UT.

Altogether, our findings suggest that, unlike ASCL1, NEUROD1 and POU2F3, YAP1 is not a subtype defining transcription factor in SCLC. This is consistent with a recent study in a patient cohort, which failed to identify a distinct YAP1 expressing SCLC subtype^7^. Although focal YAP1 expression has been detected in some primary SCLC samples^6^, our findings together with recent studies in SCLC xenograft and GEM models suggests that this is due to intra-tumoral heterogeneity^1,4,5,23^. In this context, classical ASCL1 driven SCLC can transition to a ‘neuroendocrine low’ phenotype, which is associated with expression of YAP1. This phenotypic plasticity is a feature of *RB1* null SCLC, and the emergence of neuroendocrine low, YAP1 expressing cells has been associated with chemoresistance, activation of Notch signalling and expression of mesenchymal and inflammation associated genes^1,5,23^.

SMARCA4-deficiency has been shown to be a strong predictor of sensitivity to CDK4/6 inhibitors in NSCLC and SSCOHT tumour models^24,25^. Intriguingly, several of the SCLC-Y lines that we now identify to be SMARCA4-deficient malignancies, have previously been shown to be sensitive to CDK4/6 inhibitors^9^, highlighting the clinical importance of resolving the identity of these tumours. These SMARCA4-deficient SCLC-Y cell lines may therefore provide pre-clinical models of SMARCA4-UT to accelerate the discovery of new therapeutics for this aggressive malignancy.

## MATERIALS AND METHODS

### Cell culture and transplantation studies

SCLC cell lines were obtained from American Type Culture Collection (ATCC) and cultured according to the manufacturer’s recommendations with routine testing for mycoplasma. Tumour xenografts were generated by injection of 1 x 10^6^ cells in 50% growth factor reduced Matrigel (BD Biosciences) into the flanks of CBA.Nude mice. Animal experiments were conducted according to the regulatory standards approved by the Walter and Eliza Hall Institute Animal Ethics Committee (AEC 2020.013).

Tumours (200-1000 mm^3^) were harvested, fixed in 10% neutral buffered formalin at room temperature or 4% (w/v) paraformaldehyde and paraffin embedded. Sections were stained with haematoxylin and eosin (H&E) and immunostained as described in **Supplementary Table 6**.

### Multi-omics analysis of cell line and patient samples

SCLC class was determined by CCLE RNA-seq data for driver TFs *ASCL1, NEUROD1, POU2F3*, and *YAP*. For each cell line, the TF with the highest expression among the four is used to assign the TF class. Mutations present in over 50% of cell lines within each molecular subtype were extracted via cBioportal.org^26,27^. Known pathogenic mutations were annotated using OncoKB™ and https://cancerhotspots.org available through cBioportal.org.

We used the following datasets from UT Lung SPORE: Microarray, RNA-seq gene expression data and mutation data (dbGAP Study Acession: phs001823.v1.p1). We used the following datasets from DepMap^12^: RNA-seq(CCLE_depMap_19Q1_TPM.csv), microarray (CCLE_Expression_Entrez_2012-09-29.gct), proteomics(Table_S2_Protein_Quant_Normalized.xlsx)^28^, mutation (downloaded as CCLE_DepMap_18q3_maf_20180718.txt), and structural variation dataset (CCLE_translocations_SvABA_20181221.xlsx). For each cell line, we computed the neuroendocrine score as described before^29^ but with the updated NE signature based on RNA-seq data^10^. Le Loarer datasets^15^ were downloaded from SRA under accession SRP052896 and processed with RNA-seq pipeline (https://git.biohpc.swmed.edu/BICF/Astrocyte/rnaseq) from UTSW Bioinformatics Core Facility. The CCLE for cell lines annotated as SCLC, NSCLC, or SCCOHT was merged with the Le Loarer RNA-seq data and were quantile normalized.

## ACKNOWLEDGEMENTS

We thank E.Tsui and the WEHI Histology Facility for providing immunohistochemical expertise. We are grateful to D. Boyd, L. Scott, and R. Monaghan for technical support and animal husbandry; and M. Papari-Zareei and V. Stasny for aid in searching for the original biopsies for each cell line and preservation of SCLC lines. K.D.S. is supported by the Peter and Julie Alston Centenary Fellowship and an Australian National Health and Medical Research Council (NHMRC) Project Grant 1159955. M.L.B is supported by a Snow Fellowship from the Snow Medical Research Foundation, NHMRC Investigator Grant 1196598 and Project Grant 1164054. K.D.S. and M.L.B. are supported by NHMRC Synergy Grant 2010275. JDM, LG, LC are supported by P50 CA070907, JDM by CA21338, CA213274. The bioinformatic analysis was funded by by the Cancer Prevention and Research Institute of Texas (RP150596) and UTSW ACS-IRG (IRG-21-142-16). This work was made possible through the Victorian Government Operational Infrastructure Support and Australian Government.

## SUPPLEMENTARY FIGURES

**Supplementary Figure 1.**
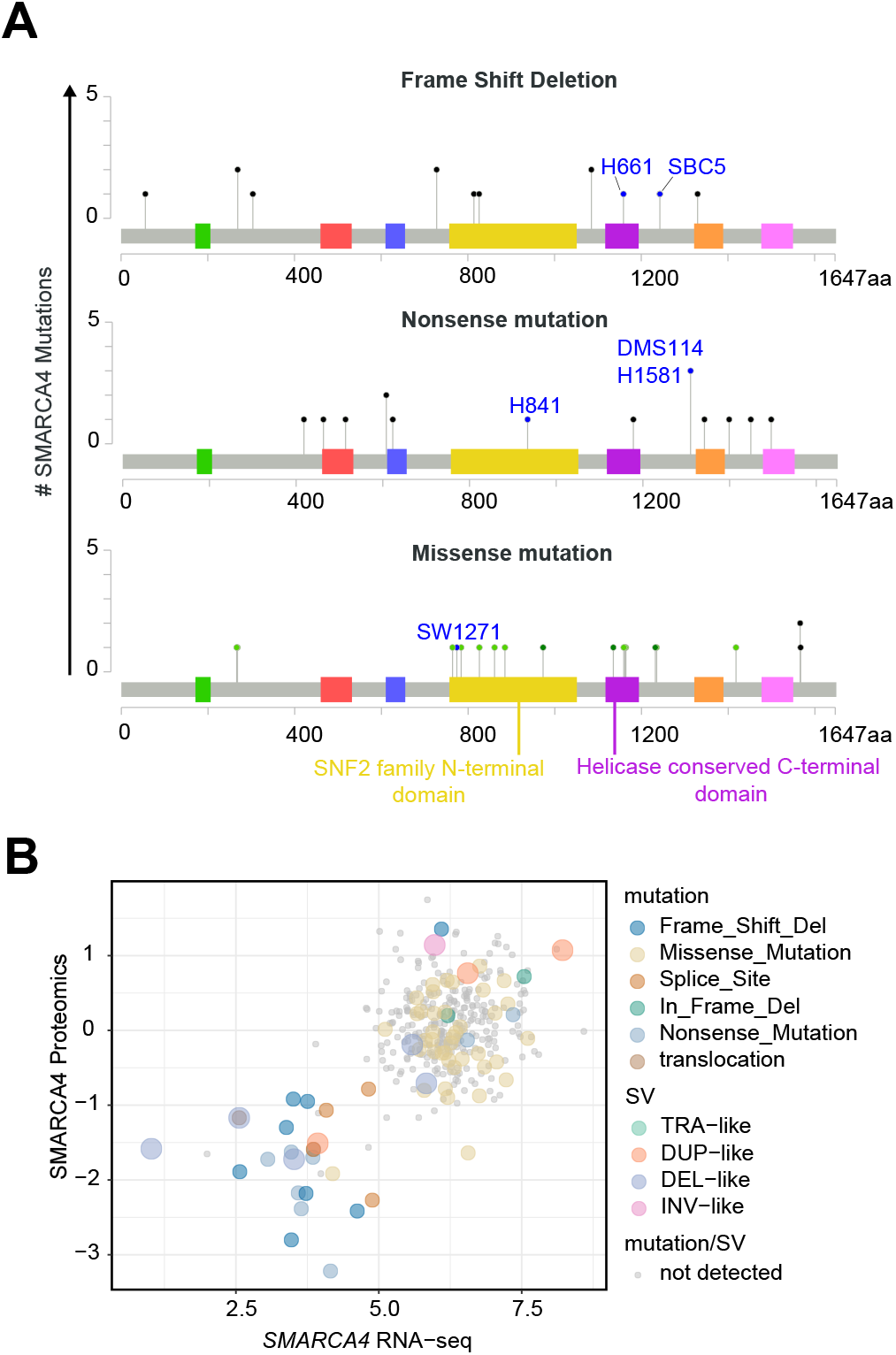
Characterisation of SMARCA4 mutations in lung cancer cell lines from the CCLE. (***A***) Lollipop diagram showing the position of frame shift deletions, nonsense and missense mutations in *SMARCA4* across lung cancer cell lines. SMARCA4-mutant SCLC-Y cell lines are annotated in blue. Mutations around the SNF2 family N-terminus and Helicase conserved C-terminal domains are important ATP binding sites for chromatin remodelling. (***B***) Key mutations associated with a decrease in SMARCA4 protein and mRNA include frame shift deletions, nonsense mutations, deletion-like structural variations and splicesite mutations. Most missense mutations do not result in a decrease in SMARCA4 protein nor mRNA.

**Supplementary Figure 2.**
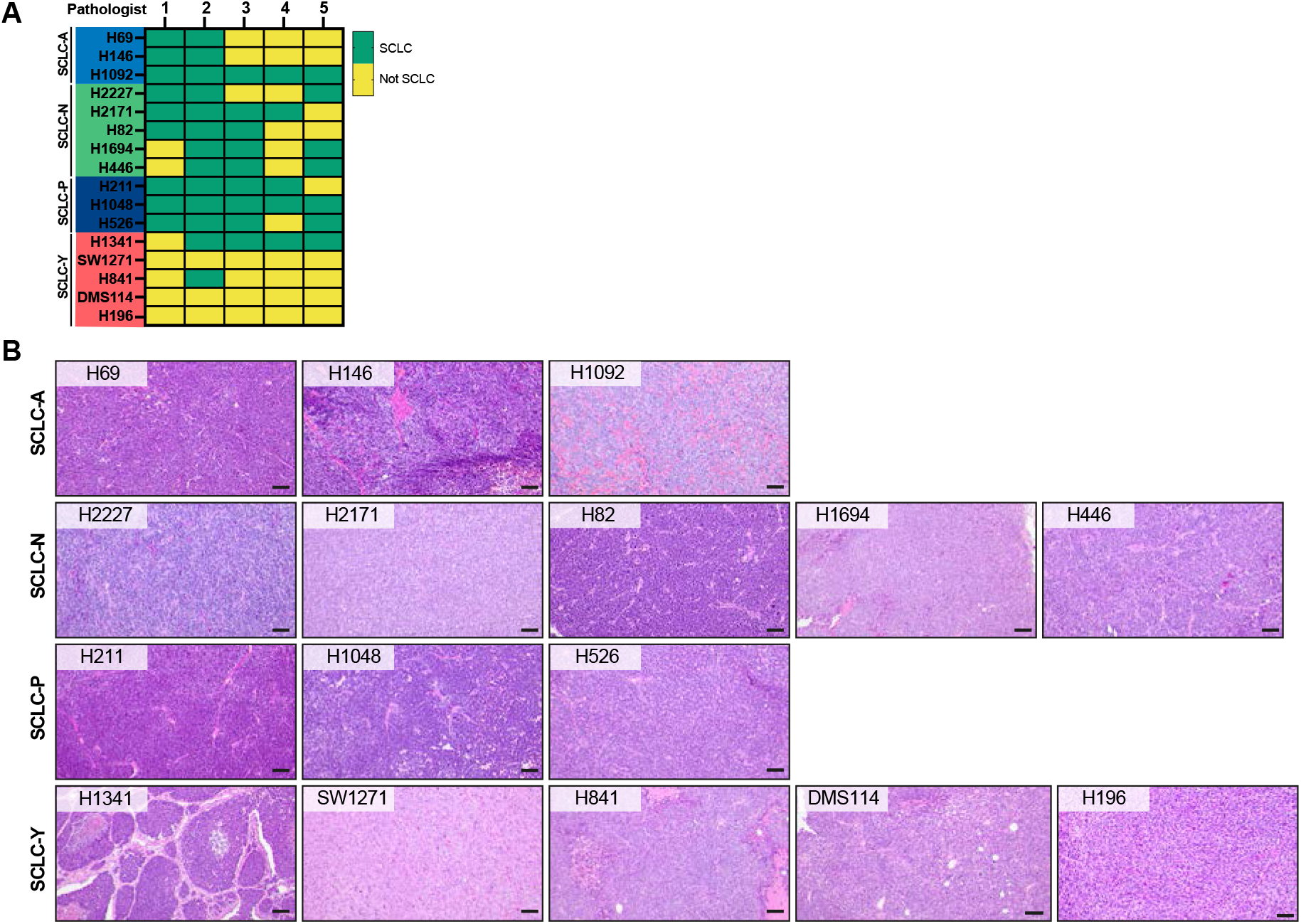
Initial histopathological evaluation of SCLC xenografts. (***A***) Heatmap of initial diagnosis (based on H&E) of SCLC cell line xenografts (row) and each pathologists’ classification (column). There was a consensus amongst the pathologists that the H&E appearance of four of the five SCLC-Y xenografts was not consistent with SCLC. (***B***) H&E images of SCLC cell line xenografts used in this study. Scale bar = 100 μm.

**Supplementary Figure 3.**
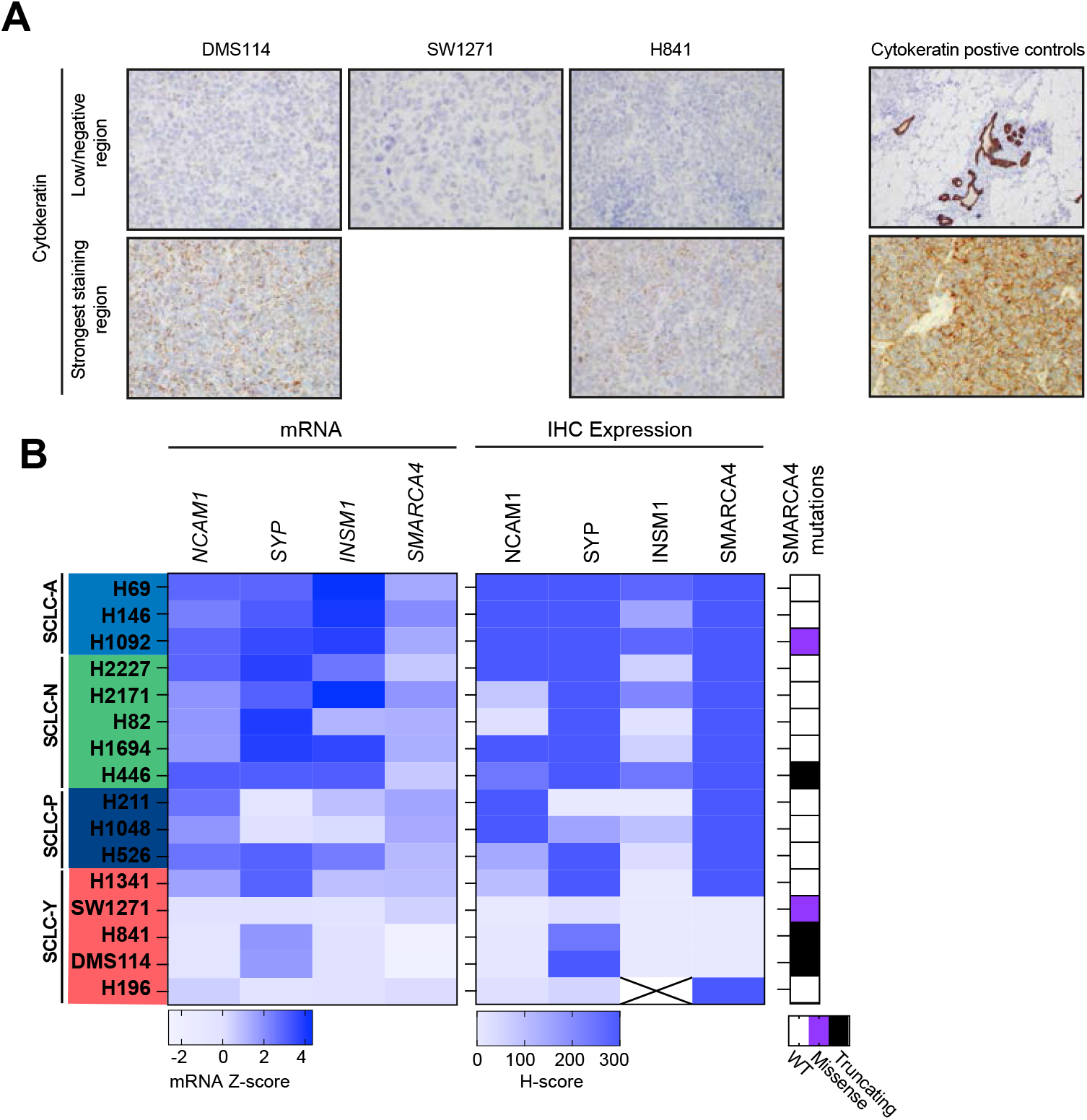
Concordance between mRNA levels and immunohistochemical staining of diagnostic neuroendocrine markers and SMARCA4 in SCLC xenografts. (***A***) Cytokeratin staining in SMARCA4-deficient SCLC-Y cell lines DMS114, SW1271 and H841 showing weak cytokeratin staining relative to the cytokeratin positive controls (right panel). (***B***) mRNA expression of neuroendocrine markers (*NCAM1, SYP, INSM1*) and *SMARCA4* compared protein levels evaluated by IHC. The INSM1 immunophenotyping in H196 was not interpretable due to a high level of non-specific background staining.

**Supplementary Figure 4.**
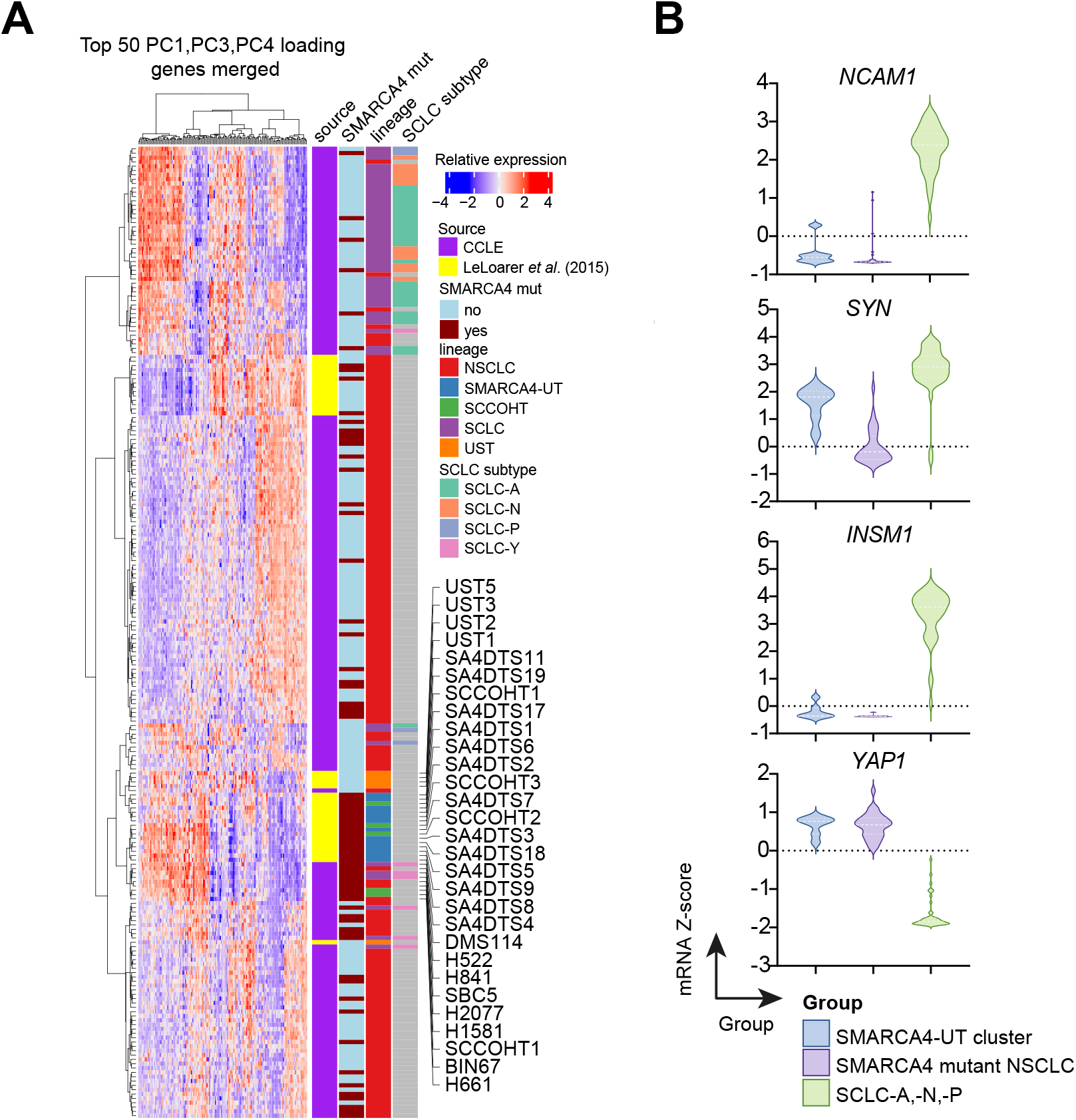
Principal component analysis of CCLE and tumour samples. (***A***) Using the top 50 genes with high loadings from PC1, PC3, and PC4, we created a smaller gene set (n=150) to cluster samples from CCLE and primary tumours from Le Loarer *et al*.^15^. Unsupervised hierarchal clustering showed a similar clustering pattern to the heatmap in Fig 3A, confirming a closer relationship of SCLC-Y lines (DMS114, H841 and SBC5) with SMARCA4-UT than SCLC. (***B***) Gene expression of neuroendocrine markers (NCAM1, synaptophysin and INSM1) and YAP1 across cell lines in the SMARCA4-UT cluster, SMARCA4-mutant NSCLC and SCLC-A, -N, -P.

**Supplementary Figure 5.**
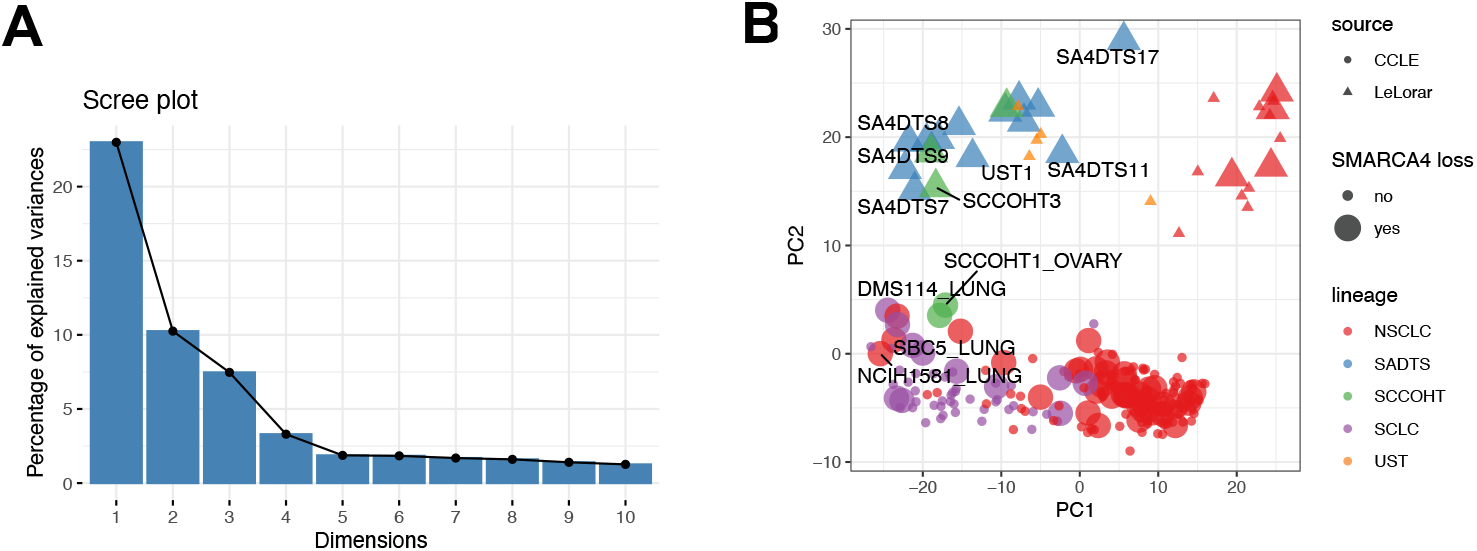
Identifying batch effects within principal component dimensions. **(A)** Based on the Scree plot, we examined the top four principal components (PCs) dimensions, which together accounts for 44% of the total variance. (***B***) Samples from CCLE and Le Loarer *et al.^15^* separate on PC2, suggesting genes correlated with PC2 is associated with batch effects between cell line and bulk-sequenced tumour samples. SMARCA4 loss was determined by integrating genetic aberration and RNA/protein expression data. For the expression data, we used model-based clustering implemented by R package *mclust^1^* to determine SMARCA4-high and SMARCA4-low cell lines. For proteomics data we used the feature for “sp|P51532|SMCA4_HUMAN”. We then merged the two sets of SMARCA4 loss status data. A loss status of “yes” is assigned to a cell line with any mutation or low expression found in any of the datasets, a loss status of “no” is assigned to a cell line with no mutation found and not a SMARCA4-low line based on any of the expression data. Abbreviations: NSCLC = Non-small cell lung cancer; SADTS = SMARCA4-deficient thoracic sarcoma (SMARCA4-UT); SCCOHT = Small cell carcinoma of the ovary hypercalcaemic type; UST = Unclassified thoracic sarcoma.

